# Urbanization impacts apex predator gene flow but not genetic diversity across an urban-rural divide

**DOI:** 10.1101/679720

**Authors:** DR Trumbo, PE Salerno, KA Logan, M Alldredge, RB Gagne, CP Kozakiewicz, S Kraberger, N Fountain-Jones, ME Craft, S Carver, HB Ernest, K Crooks, S VandeWoude, WC Funk

**Affiliations:** Department of Biology, Colorado State University, Fort Collins, CO 80523 USA; Colorado Parks and Wildlife, Montrose, CO 81401 USA; Colorado Parks and Wildlife, Fort Collins, CO 80526 USA; Department of Microbiology, Immunology, and Pathology, Colorado State University, Fort Collins, CO 80523 USA; Department of Biological Sciences, University of Tasmania, Hobart, TAS 7005 Australia; Department of Veterinary Population Medicine, University of Minnesota, Saint Paul, MN 55108 USA; Department of Veterinary Sciences, University of Wyoming, Laramie, WY 82070 USA; Department of Fish, Wildlife, and Conservation Biology, Colorado State University, Fort Collins, CO 80523 USA; Graduate Degree Program in Ecology, Colorado State University, Fort Collins, CO 80523 USA

**Keywords:** landscape genomics, gene flow, genetic diversity, effective population size, urbanization, *Puma concolor*

## Abstract

Apex predators are important indicators of intact natural ecosystems. They are also sensitive to urbanization because they require broad home ranges and extensive contiguous habitat to support their prey base. Pumas (*Puma concolor*) can persist near human developed areas, but urbanization may be detrimental to their movement ecology, population structure, and genetic diversity. To investigate potential effects of urbanization in population connectivity of pumas, we performed a landscape genomics study of 134 pumas on the rural Western Slope and more urbanized Front Range of Colorado, USA. Over 12,000 single nucleotide polymorphisms were genotyped using double-digest, restriction site-associated DNA sequencing (ddRADseq). We investigated patterns of gene flow and genetic diversity, and tested for correlations between key landscape variables and genetic distance to assess the effects of urbanization and other landscape factors on gene flow. Levels of genetic diversity were similar for the Western Slope and Front Range, but effective population sizes were smaller, genetic distances were higher, and there was more overall population substructure in the more urbanized Front Range. Forest cover was strongly positively associated with puma gene flow on the Western Slope, while impervious surfaces restricted gene flow and more open, natural habitats enhanced gene flow on the Front Range. Landscape genomic analyses revealed differences in puma movement and gene flow patterns in rural versus urban settings. Our results highlight the utility of dense, genome-scale markers to document subtle impacts of urbanization on a wide-ranging carnivore living near a large urban center.

## Introduction

Urbanization is a major threat to biodiversity, and in particular to apex predators with broad home ranges (Cohen 2003; Theobald 2005; Crooks *et al.* 2017). Habitat fragmentation due to urbanization can have important impacts on predator movement, disease, and survival (Markovchick-Nicholls *et al* 2008; Carver *et al.* 2016; Fountain-Jones *et al.* 2017). This reduced connectivity can lead to smaller, more isolated populations, where less gene flow and genetic diversity, as well as smaller effective population sizes (Riley *et al.* 2006; Vandergast *et al.* 2007; Ernest *et al.* 2014) ultimately cause local and regional extirpations through environmental and demographic stochasticity and inbreeding depression (Allendorf *et al.* 2013). Moreover, increased human recreational activities in wildlife habitats associated with nearby urbanization can change wildlife movement patterns and habitat usage, exacerbating the impacts of fragmentation (McKinney 2002; Lewis *et al.* 2015). As human populations continue to expand worldwide, urban areas are becoming larger and more extensive on the landscape. However, we do not fully understand how urbanization affects natural ecosystems near wildland-urban interfaces (Radeloff *et al.* 2005; Magle *et al.* 2012).

Large carnivores are important indicators of intact natural ecosystems, as they require an abundant and sustainable prey base, as well as high habitat connectivity to support their broad home ranges (Sergio *et al.* 2006, 2008). However, understanding the effects of urbanization on large carnivores is difficult due to their low population densities and secretive nature (Logan and Sweanor 2001; Riley *et al.* 2006; Hornocker and Negri 2009). Camera traps, radio-telemetry, and GPS collars provide valuable information on animal home ranges and population sizes (e.g., Lewis *et al.* 2015; Blecha *et al.* 2018), but these studies are expensive, time consuming, and can only monitor a small fraction of the total population for limited time periods. Population and landscape genetics can provide additional, complementary techniques for a more detailed understanding of wildlife populations (Epps *et al.* 2007; Lowe and Allendorf 2010; Balkenhol *et al.* 2016). Genetic studies provide an indicator of functional landscape connectivity through measures of gene flow, effective population sizes of breeding individuals, and cost-efficient monitoring of genetic diversity across broad geographic areas (McRae *et al.* 2005; Solberg *et al.* 2006). Moreover, recent high-throughput sequencing technologies enable the genotyping of many more thousands of loci than previously possible, providing higher power to detect the often subtle population genetic structure of wide-ranging species such as large carnivores (Luikart *et al.* 2003; Holderegger *et al.* 2006).

Pumas (*Puma concolor*; other common names include mountain lions, cougars, panthers, catamounts) are a large, apex predator with one of the broadest latitudinal ranges of any terrestrial carnivore, spanning western North America, Central America, and South America (Hornocker and Negri 2009). Pumas are sensitive to urbanization, requiring broad-scale landscape connectivity to persist, and are thus useful indicators for monitoring the effects of urban fragmentation (Beier 1995; Crooks 2002; Maletzke *et al.* 2017). Given sufficient habitat area and landscape connectivity, however, pumas can still persist within and adjacent to urban systems (Wilmers *et al.* 2013; Riley *et al.* 2014; Lewis *et al.* 2015; Zeller *et al.* 2017; Blecha *et al.* 2018). Furthermore, the substantial area requirements of large carnivores such as pumas can enhance their role as “umbrella” species, whose protection also benefits co-occurring species through broad-scale habitat preservation (Thorne *et al.* 2006).

The southern Rocky Mountains in western Colorado, USA support natural habitats with high puma densities, as well as many rural and urban human developments (Hornocker and Negri 2009). The Western Slope of the Rocky Mountains primarily consists of large areas of contiguous public wildlands with an abundant prey base for pumas, interspersed with small rural and exurban developments, including the Uncompahgre Plateau region near the town of Montrose (Western Slope Study Area; Figure 1). In contrast, the Front Range is a rapidly urbanizing, major metropolitan area on the Eastern Slope of the Continental Divide, where urbanization is spreading from lower elevation areas in and around the Denver Metropolitan Area into adjacent wildland habitats in the foothills of the Rocky Mountains. Pumas continue to persist near this wildland-urban interface, including adjacent to the city of Boulder on the western edge of the Denver Metropolitan Area (Front Range Study Area; Figure 1; Lewis *et al.* 2015; Moss *et al.* 2016a). From 2010 – 2017, Colorado was the 8th fastest growing U.S. state by population (577,829 residents added) and the 6th fastest by percentage (11.5% population growth; U.S. Census Bureau 2017), with most of this growth occurring along the eastern edge of the Front Range. Thus, comparative studies of puma movement and gene flow in one of the most populous states in the mid-continental USA, which also supports a robust puma population, can provide insight into the effects of urbanization on this important apex predator.

**Figure 1:**
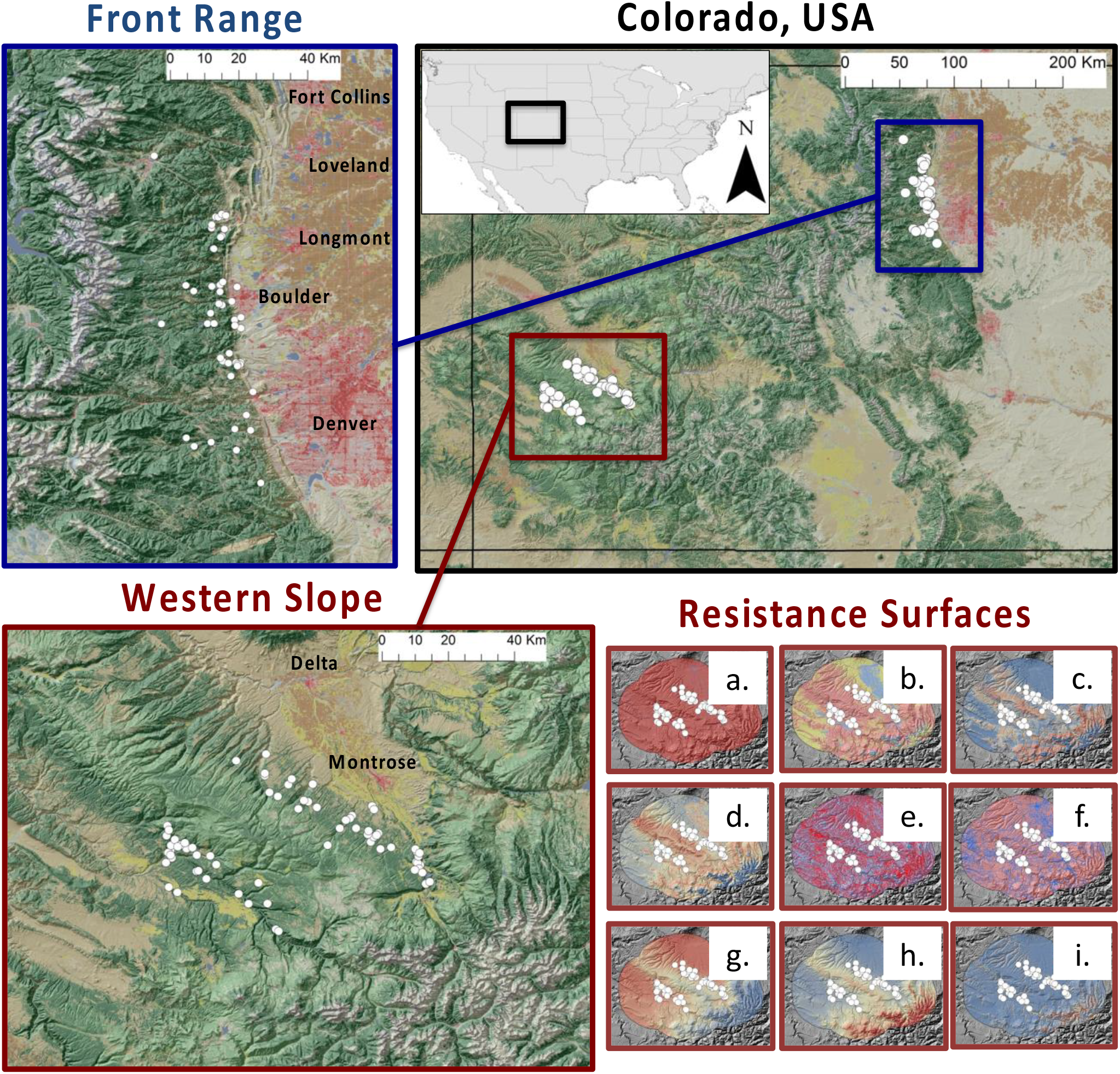
Study area in the Western Slope and Front Range of the southern Rocky Mountains of Colorado, USA. Landscape genomic analyses included 78 pumas from the Western Slope and 56 pumas from the Front Range (white circles). Resistance surfaces, shown for the Western Slope, represent alternative hypotheses of the effects of landscape variables on puma dispersal and gene flow (red=high gene flow, blue=low gene flow) for: (a) percent impervious surface cover (negative effect on gene flow),(b) land cover (forested, open-natural, and developed: positive, neutral, and negative effects on gene flow), (c) percent tree canopy cover (positive effect), (d) vegetation density (positive effect), (e) river and stream riparian corridors (positive effect), (f) roads (negative effect), (g) minimum temperature of the coldest month (negative effect), (h) annual precipitation (positive effect), and (i) topographic roughness (positive effect). We also tested isolation by geographic Euclidean distance. Land cover base maps show forests (green), shrub and grasslands (tan), urban areas (red), agriculture and ranchlands (brown and yellow), and alpine tundra (grey).

Here, we tested how different landscape factors, including urbanization, enhance or restrict gene flow and genetic diversity in a large apex predator across an urban-rural divide in Colorado, USA. A large sample (n = 134) of pumas were utilized from (a) the rural Western Slope and (b) the more urbanized Front Range (Figure 1). We used double digest restriction site associated DNA sequencing (ddRADseq) to genotype pumas at 12,444 single nucleotide polymorphism (SNP) loci to evaluate the potential differences in gene flow, effective population sizes, genetic diversity, and population structure in these two different landscapes. We tested landscape genomic hypotheses by correlating key landscape factors with puma genetic distance measures. We hypothesized that pumas in the more urbanized Front Range would have smaller effective population sizes, (b) lower levels of genetic diversity, and (c) more landscape factors related to urbanization that restrict gene flow, relative to the rural Western Slope landscape.

## Materials and Methods

### Samples and sequences

Puma blood and tissue samples were collected as part of ongoing monitoring efforts by Colorado Parks and Wildlife in both the Western Slope and Front Range regions of the southern Rocky Mountains of Colorado, USA (Figure 1; Lewis *et al.* 2015; Carver *et al.* 2016). Samples were collected from 2005-2014 on the Western Slope and 2007-2013 on the Front Range. Western Slope samples consisted of 36 males and 42 females, and Front Range samples consisted of 24 males, 31 females, and 1 puma of unknown sex. Our sampling represents a large proportion of the resident pumas present in both regions during the sampling period, as Lewis *et al.* (2015) estimated 14.4 (S.E. 1.6) and 14.7 (S.E. 1.3) resident pumas occupying the Western Slope and Front Range study areas at a single time point, respectively, from motion camera and telemetry data collected in 2009 and 2010.

Genomic DNA was extracted from tissue or blood using QIAGEN DNeasy Blood & Tissue kits (QIAGEN Inc., Valencia, CA). We genotyped a total of 78 individuals from the Western Slope and 56 individuals from the Front Range using the ddRADseq protocol described in Peterson *et al.* (2012) and sequenced on Illumina HiSeq 2500 and 4000 machines (Illumina, San Diego, California) using 100bp single-end sequencing at the University of Oregon Genomics Facility (https://gc3f.uoregon.edu). We tested 9 different combinations of restriction enzymes on puma samples for digestion efficiency and evaluated the size ranges of fragment distributions using an Agilent Tapestation 2200 (Agilent Genomics, Santa Clara, California). We chose the digest enzymes EcoRI-HF (6bp recognition) and NlaIII (4bp recognition) and a target fragment size range of 300–400 bp (excluding adapters). We used a Blue Pippin with a 2%, internal standard, 100-600 bp gel cartridge (Sage Science, Beverly, Massachusetts) for size selection and a biotinylated P2 adapter with DynaBeads^®^ (Peterson *et al.* 2012) to purify the polymerase chain reaction (PCR) template for the final enrichment. PCR was performed for 12 cycles and five reactions were tested for each pool of individuals. We initially genotyped 16 individuals multiplexed into an Illumina 2500 HiSeq lane to estimate maximum multiplexing based on a target of >12X coverage per locus. After assessment of locus coverage, we proceeded to multiplex 48 and 70 individually-barcoded samples on Illumina 2500 and 4000 HiSeq lanes, respectively, using the Peterson *et al.* (2012) flex adaptors.

### Bioinformatics pipeline and filters

We evaluated read quality for each sequencing lane using FastQC (bioinformatics.babraham.ac.uk) and assembled our SNP dataset *de novo* using Stacks v 1.41 (Catchen *et al.* 2013). Details on Stacks code and parameter settings used are on the GitHub repository; github.com/pesalerno/PUMAgenomics. We demultiplexed and filtered sequencing reads using the program *process_radtags* in Stacks. Due to sensitivity of downstream genotyping with different Stacks parameter settings (Mastretta-Yanes *et al.* 2015; Paris *et al.* 2017), we incorporated individual sample replicates in library preparations. In each library, we included 3 within and 3 between library replicates, which were used for estimating genotyping error rates for different combinations of parameters used to construct loci with the *denovo_map.pl* Stacks pipeline. We ran 11 different *de novo* assemblies varying 4 different Stacks parameters that affect locus, allele, and SNP error rates and the number of loci genotyped, consisting of (1) minimum number of identical, raw reads required to create a stack (−m), (2) number of mismatches allowed between loci when processing a single individual (−m), (3) number of mismatches allowed between loci when building the catalog (−n), and (4) maximum number of stacks at a single de novo locus (-max_locus_stacks) (Table S1; Mastretta-Yanes *et al.* 2015). Locus error rate was calculated as the number of loci present in only one of the samples of a replicate pair divided by the total number of loci, allele error rate was the number of allele mismatches between replicate pairs divided by the number of loci, and SNP error rate was the proportion of SNP mismatches between replicate pairs.

After identifying the most supported parameter settings that minimized locus, allele, and SNP error rates, while maximizing the number of SNPs (−m = 3, -M = 4, −n = 4, max_locus_stacks = 3; Table S1), we exported the SNP matrix with the *populations* program in Stacks (Catchen *et al.* 2013), retaining SNPs that were present in at least 20% of individuals by population, and retaining a single random SNP per locus. This matrix was further filtered for missing data in Plink v. 1.07, first by locus, then by individual, and then by minor allele frequency (MAF) using multiple combinations of thresholds for reducing missing data in the matrix (see github.com/pesalerno/PUMAgenomics). After evaluating missing data from SNP matrices, we retained the matrix with a more stringent locus filter (excluding loci missing >25% individuals) and a less stringent filter on minor allele frequency (excluding loci with MAF < 0.01). We additionally filtered loci that were found at position 95 (the last position of our reads) due to a higher number of SNPs present in this position, suggesting increased error rates due to low sequence quality towards the end of the sequencing read. In order to compare landscape resistances with putatively neutral loci, we used a Principal Components Analysis (PCA) to identify loci showing strong signatures of selection relative to neutral background genomic variation with the program PCAdapt (Luu *et al.* 2016). We found twelve, putatively adaptive, outlier loci using a false discovery rate of 10%, so we filtered these outliers out for downstream landscape genomic analyses to avoid confounding neutral demographic patterns with patterns generated by loci under selection.

### Population genomics and structure

Population genomic statistics were calculated for the two sampling regions, the Western Slope and Front Range (Figure 1). Observed and expected heterozygosity (H_obs_ and H_exp_), nucleotide diversity (π), inbreeding coefficient (F_IS_), and population genetic differentiation (F_ST_) were calculated using the *populations* program in Stacks with SNP loci that passed previous filters, excluding a single individual (sample_1382) that did not pass the 75% missing data threshold. We estimated allelic richness (A_r_) using HP-RARE 1.0 (Kalinowski 2005), which corrects for variance in sample sizes using rarefaction. Two complementary, individual-based genetic distances were calculated: proportion of shared alleles distance (D_ps_; Bowcok *et al.* 1994) using the adegenet R v. 3.3.3 package and relatedness distance (r; Smouse and Peakall 1999) using the PopGenReport R package. We then calculated mean genetic distance among individuals for each region, corrected for geographic distance (i.e., genetic distance per km), since individuals that are farther apart are expected to have higher genetic distances due to neutral isolation by distance population processes (Wright 1942; Balkenhol *et al.* 2016). Effective population sizes (*N*_*e*_) were estimated using the linkage disequilibrium method in NeEstimator v. 2.01 (Do *et al.* 2014), using the correction for chromosome number (Waples *et al.* 2016), which has been shown to be a robust method for inferring *N*_*e*_ using SNP datasets and large sample sizes (Waples 2016; Waples *et al.* 2016). We evaluated overall genetic structure as well as genetic differentiation among the two sampling sites (Western Slope and Front Range) using PCA and Discriminant Analysis of Principal Components (DAPC) in the R package adegenet (Jombart 2008) and Admixture ancestry analysis (Alexander *et al*. 2009). We used the function assignplot to identify individuals that were putative migrants or admixed based on the individual DAPC assignment probabilities. We used the find.clusters command in adegenet and minimized cross validation error in Admixture to estimate the number of populations (i.e., K).

### Landscape genomics

Geographic Information Systems (GIS) data were collected for different landscape factors that we hypothesized would affect puma dispersal and gene flow in Colorado. Table 1 provides details on GIS data sources, spatial resolution, and ecological justification for each landscape factor. Study area extents were calculated and landscape variables were compared across regions by buffering individual data points by a typical female puma dispersal distance of 34.6 km (Logan and Sweanor 2001), dissolving overlapping buffers, and calculating zonal statistics within each region (Western Slope and Front Range) using ArcGIS v. 10.1 (ESRI, Redlands, California). Landscape data were converted into resistance surfaces using the Reclassify and Raster Calculator tools in ArcGIS. The following hypothesized relationships of landscape factors with puma gene flow were modeled: percent impervious surface cover (negative effect on gene flow), land cover (forested, open-natural, and developed: positive, neutral, and negative effects on gene flow, respectively), percent tree canopy cover (positive effect), vegetation density (positive effect), river and stream riparian corridors (positive effect), roads (negative effect), minimum temperature of the coldest month (negative effect), annual precipitation (positive effect), topographic roughness (positive effect), and elevation (negative effect). Additionally, we included an isolation by geographic distance model, which would be supported if none of the landscape variables had an effect on gene flow except for straight line, Euclidean distance between individuals (Wright 1942; Balkenhol *et al.* 2016). Table S2 describes methods and justification for converting raw landscape variables to resistance surfaces.

**Table 1:**
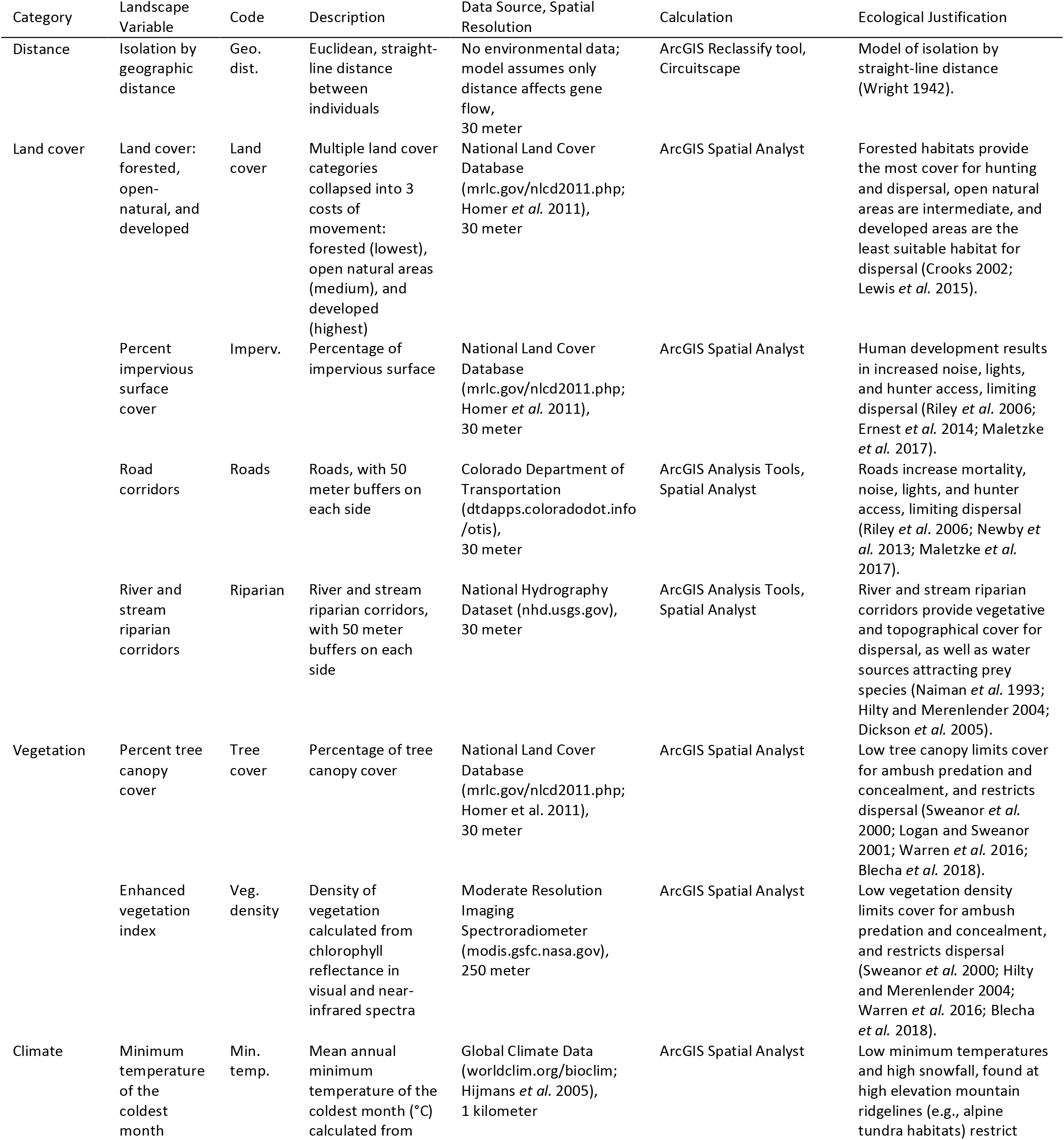
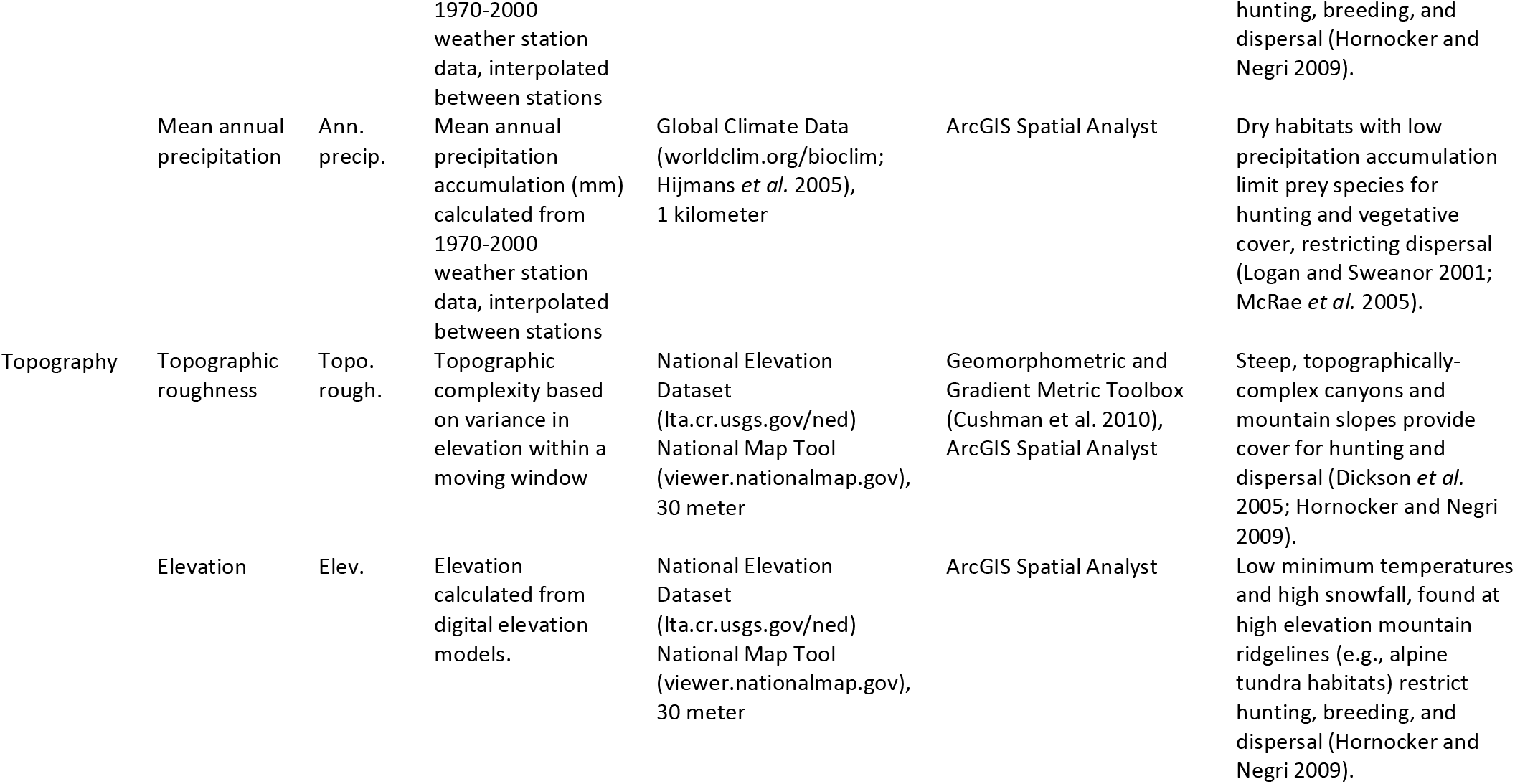
Environmental variables used for landscape genomic analyses, data sources, spatial resolution, and ecological justification.

Two genetic distance measures were used as response variables in landscape genomic analyses: proportion of shared alleles distance (D_ps_; Bowcok *et al.* 1994) and relatedness distance (r; Smouse and Peakall 1999). Environmental resistances among individuals were calculated using Circuitscape (McRae 2006) for each landscape resistance surface (McRae 2006; Row *et al.* 2017). Circuitscape resistances are a useful tool in landscape genetics because they summarize all potential movement pathways simultaneously, as opposed to least cost paths that evaluate only a single idealized pathway, and thus assume the study organism has complete knowledge of the landscape and always chooses the ideal pathway (McRae 2006; Balkenhol *et al.* 2015). Landscape variables were tested for multicollinearity, both prior to and after calculating environmental resistances in Circuitscape, to ensure Pearson’s r correlations < 0.7 and variance inflation factor (VIF) scores < 5 in final landscape genomics models, as collinearity can cause instability in parameter estimation in regression models (Tables S3 and S4; Warren *et al.* 2010; Dormann *et al.* 2012; Rowe *et al.* 2017).

Two complementary methods were used to estimate the effects of environmental resistances on genetic distances: multiple regression on distance matrices (MRDM; Legendre *et al.* 1994) using PERMUTE v.3.4 and maximum likelihood of population effects (MLPE; Clark *et al.* 2002; van Strien *et al.* 2012; Row *et al.* 2017) using the lme4 R package. MRDM is a permutational, distance matrix-based approach that has been traditionally used in landscape genetic analyses, whereas MLPE is a newer linear mixed effects modeling technique that models pairwise comparisons as a random effect and environmental resistances as fixed effects (Balkenhol *et al.* 2016). Recent evaluations of landscape genetic approaches found linear mixed effects modeling using MLPE to be more accurate, although both approaches performed well (Shirk *et al.* 2017). Therefore, we included the traditional MRDM approach as well as MLPE in order to utilize multiple, complementary techniques for inferring associations between landscape features and gene flow. For MRDM and MLPE, genetic distances were the response variable and environmental resistances were explanatory variables. Additionally for MLPE, a random effect matrix of individual comparisons was included to control for the non-independent, pairwise structure of the data, and landscape resistances were standardized to units of standard deviation centered on the mean (van Strien *et al.* 2012; Row *et al.* 2017). Models were ranked using the Bayesian information criterion (BIC), and top models within 5 BIC units are reported (Richards 2015).

## Results

### Genotyping and filtering SNP matrices

Initial Stacks processing retained a single random SNP per 95 bp read and SNPs present in at least 20% of individuals by population, resulting in a matrix of 98,813 SNPs. These SNPs were further filtered in Plink by removing loci that were present in less than 75% of individuals, which resulted in a matrix of 20,355 SNPs. Only a single individual was excluded based on our >75% missing loci per individual threshold. After excluding SNPs present in the 95^th^ sequencing base position and with minor allele frequency <0.01, we retained 12,456 SNPs. PCAdapt detected twelve outlier loci, putatively under selection, while accounting for population structure (K=2). After removing these putatively adaptive loci, the final neutral dataset contained 12,444 SNPs (Table S1; github.com/pesalerno/PUMAgenomics).

### Population genomics and structure

The two study areas encompass similar geographic extents: 11,889 km^2^ for the Western Slope and 11,958 km^2^ for the Front Range (Table 2). Measures of genetic diversity (H_obs_, H_exp_, π, A_r_,) and inbreeding (F_IS_) were similar for the Western Slope and Front Range (Table 2). However, the effective population size (*N*_*e*_) was smaller, mean genetic distances among individuals (D_PS_/km and r/km) were higher, and there was more overall population substructure in the more urbanized Front Range (Table 2, Figure 2). We also calculated *N*_*e*_ using subsets of individuals (i.e., pre and post-2010 individuals in the Front Range, pre and post-2011 individuals in the Western Slope), since multiple overlapping generations may bias effective population size estimates low or high (Waples 2016; Waples et al. 2016). *N*_*e*_ remained consistently higher in the Western Slope, although it differed between the earlier and later sampling periods there, and indicated the population may be expanding (Table S5). We found a detectable signature of population differentiation between the Western Slope and Front Range regions based on a PCA and DAPC, and Admixture ancestry analysis indicated K=2 was the best supported value of K by minimizing cross validation error (Figure 2; Alexander *et al*. 2009). The proportion of correct individual assignment to populations based on DAPC (Figure 2b), which attempts to minimize within population distances and maximize between population distances (Jombart 2008), was high for most individuals in both the Western Slope (0.98) and the Front Range (0.96). However, the DAPC assignplot also identified admixed individuals and putative migrants between regions, including a female and a male in the Front Range that assigned mostly to the Western Slope, and an admixed male in the Western Slope that assigned mostly to the Front Range (Figure 2b). We also analyzed both regions separately for population substructure (Figure S1), and there was no signature of population differentiation within the Western Slope or Front Range, further supporting two populations.

**Table 2:** Study areas (km^2^), number of individuals genotyped (N_gen_), and population genomic parameter estimates from the Western Slope and Front Range of Colorado. Population genomic measures are observed heterozygosity (H_obs_), expected heterozygosity (H_exp_), nucleotide diversity (π), allelic richness (A_r_), inbreeding coefficient (F_IS_), genetic differentiation among populations (pairwise F_ST_), mean genetic distance among individuals corrected for geographic distance (D_PS_ and r per km) with standard errors (S.E.), and effective population size (N_e_) with 95% confidence intervals (C.I.) based on parametric bootstrapping.

| Region | Area (km<sup>2</sup>) | N<sub>gen</sub> | H<sub>obs</sub> | H<sub>exp</sub> | $\pi$ | A<sub>r</sub> | F<sub>IS</sub> | F<sub>ST</sub> | D<sub>PS</sub>/km (S.E.) | r/km (S.E.) | N<sub>e</sub> (95% C.I.) |
| --- | --- | --- | --- | --- | --- | --- | --- | --- | --- | --- | --- |
| Western Slope | 11889 | 78 indiv. | 0.240 | 0.272 | 0.0029 | 1.93 | 0.117 | 0.024 | 0.28 (0.05) | 0.15 (0.03) | 69.3 (66.2-72.4) |
| Front Range | 11958 | 56 indiv. | 0.242 | 0.263 | 0.0028 | 1.89 | 0.084 |  | 0.46 (0.16) | 0.24 (0.09) | 40.2 (38.7-41.7) |

**Figure 2:**
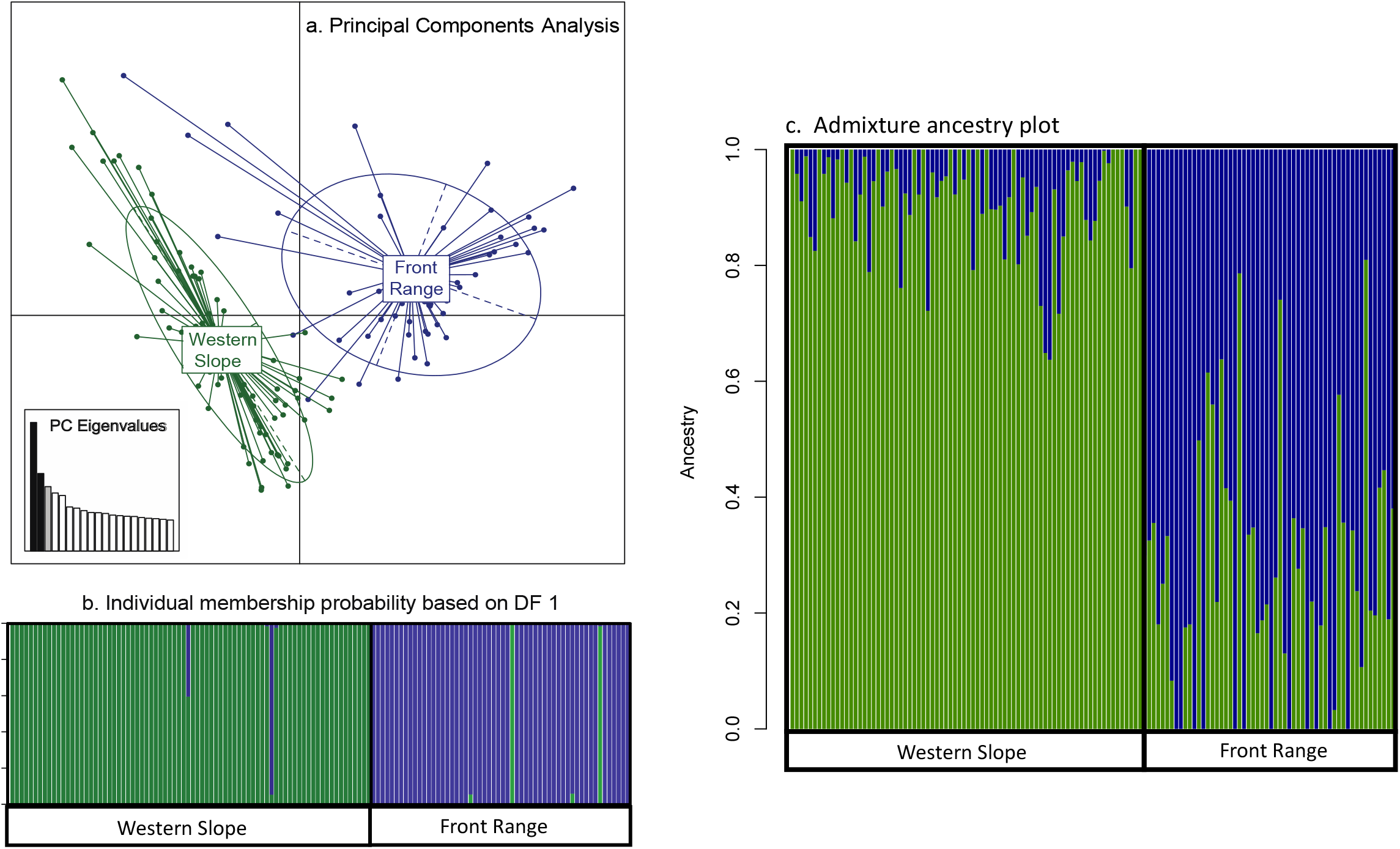
Population structure from (a) Principal Components Analysis (PCA), (b) Discriminant Analysis of Principal Components (DAPC), and (c) Admixture analysis. Individuals assigned to the Western Slope and Front Range are green and blue, respectively. K=2 was most supported in Admixture analysis using cross validation error.

### Landscape Genomics

The Front Range has more urban development than the Western Slope, with more impervious surface cover and a higher density of roads (Figure 1, Table 3, Table S2). The Front Range also has more tree canopy cover, higher vegetation density, and higher annual precipitation than the Western Slope (Table 3), likely due to the high desert habitats (i.e., the Colorado Plateau ecoregion) in the Western Slope being drier than the grassland and shrub habitats found at lower elevations of the Front Range (i.e., the Great Plains ecoregion; McMahon *et al.* 2001).

**Table 3:**
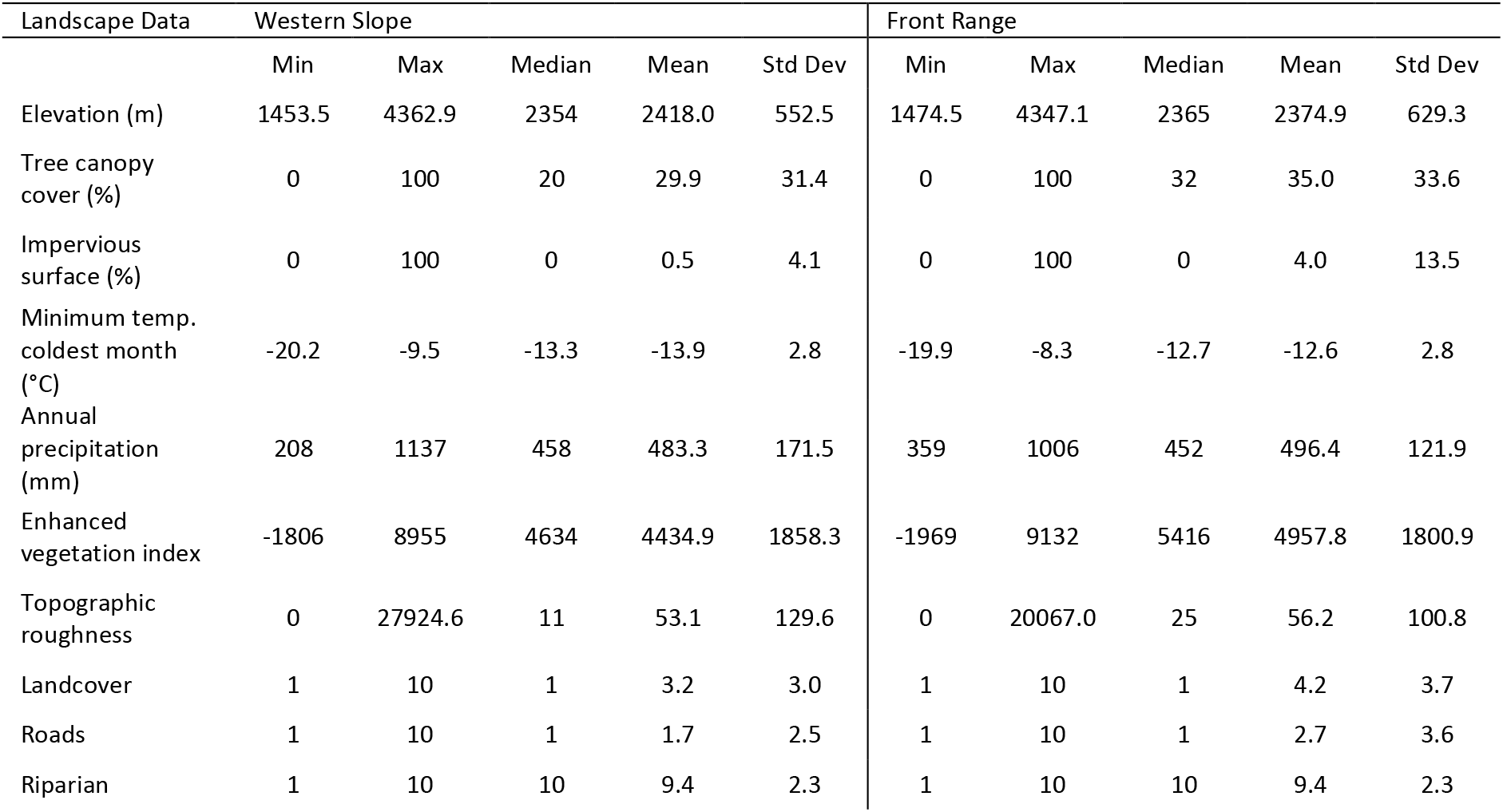
Habitat differences between the Western Slope and Front Range of Colorado. Units are percent cover for impervious surface and tree canopy cover; resistance values for land cover, river and stream riparian corridors, and roads; degrees Celsius for temperature; millimeters for precipitation; meters for elevation; and unitless measurements based on chlorophyll reflectance and variance in elevation, respectively, for enhanced vegetation index and topographic roughness.

| Landscape Data | Western Slope |  |  |  |  | Front Range |  |  |  |  |
| --- | --- | --- | --- | --- | --- | --- | --- | --- | --- | --- |
|  | Min | Max | Median | Mean | Std Dev | Min | Max | Median | Mean | Std Dev |
| Elevation (m) | 1453.5 | 4362.9 | 2354 | 2418.0 | 552.5 | 1474.5 | 4347.1 | 2365 | 2374.9 | 629.3 |
| Tree canopy cover (%) | 0 | 100 | 20 | 29.9 | 31.4 | 0 | 100 | 32 | 35.0 | 33.6 |
| Impervious surface (%) | 0 | 100 | 0 | 0.5 | 4.1 | 0 | 100 | 0 | 4.0 | 13.5 |
| Minimum temp. coldest month (°C) | -20.2 | -9.5 | -13.3 | -13.9 | 2.8 | -19.9 | -8.3 | -12.7 | -12.6 | 2.8 |
| Annual precipitation (mm) | 208 | 1137 | 458 | 483.3 | 171.5 | 359 | 1006 | 452 | 496.4 | 121.9 |
| Enhanced vegetation index | -1806 | 8955 | 4634 | 4434.9 | 1858.3 | -1969 | 9132 | 5416 | 4957.8 | 1800.9 |
| Topographic roughness | 0 | 27924.6 | 11 | 53.1 | 129.6 | 0 | 20067.0 | 25 | 56.2 | 100.8 |
| Landcover | 1 | 10 | 1 | 3.2 | 3.0 | 1 | 10 | 1 | 4.2 | 3.7 |
| Roads | 1 | 10 | 1 | 1.7 | 2.5 | 1 | 10 | 1 | 2.7 | 3.6 |
| Riparian | 1 | 10 | 10 | 9.4 | 2.3 | 1 | 10 | 10 | 9.4 | 2.3 |

Prior to running Circuitscape, landscape raster surfaces were largely uncorrelated (i.e., Pearson’s r < 0.7), with the exception of elevation, which was positively correlated with annual precipitation and negatively correlated with minimum temperature of the coldest month in both regions, and vegetation density, which was negatively correlated with annual precipitation in the Front Range (Table S3). After Circuitscape analyses, environmental resistance variables showed more collinear relationships than raw raster surfaces (Table S4), likely due to Circuitscape resistances being higher for individuals separated by larger geographic distances (McRae 2006). Therefore, we removed landscape variables from both regions that were strongly correlated with many other variables, until all VIF scores were less than 10 (Row *et al.* 2017). Variables retained were geographic distance, river and stream riparian corridors, roads, impervious surface cover, tree canopy cover, vegetation density, and minimum temperature of the coldest month.

However, vegetation density was still correlated with geographic distance in both regions, and impervious surface was correlated with geographic distance and tree canopy cover in the Western Slope (Table S4). We removed these variables as well, resulting in Pearson’s r correlations less than 0.7 and VIF scores less than or equal to 4.1 and 3.5 in the Western Slope and Front Range, respectively, for all explanatory variables. Thus final MRDM and MLPE models for the Western Slope included geographic distance, tree canopy cover, stream and river riparian corridors, roads, and minimum temperature of the coldest month; and for the Front Range included the same landscape variables plus impervious surface cover.

Landscape genomic patterns of pumas were different in the rural Western Slope compared to the more urbanized Front Range, with the exception of geographic distance being supported in both regions (Tables 4 and 5). In the Western Slope, tree canopy cover was consistently positively correlated with gene flow in MRDM and MLPE models, and low minimum temperatures of the coldest month (i.e., those found in high elevation, alpine tundra habitats) were negatively correlated gene flow in one MLPE model (Tables 4 and 5). In contrast, in the Front Range, tree canopy cover and percent impervious surface cover were negatively associated with gene flow in the top MLPE models (Table 5). Since the relationship between tree cover and gene flow was the opposite of what we hypothesized in the Front Range, we also inverted the tree cover resistance surface (i.e., making higher tree cover = higher resistance), reran Circuitscape and MLPE analyses, and higher tree cover still showed significant negative correlations with gene flow in this region.

**Table 4:**
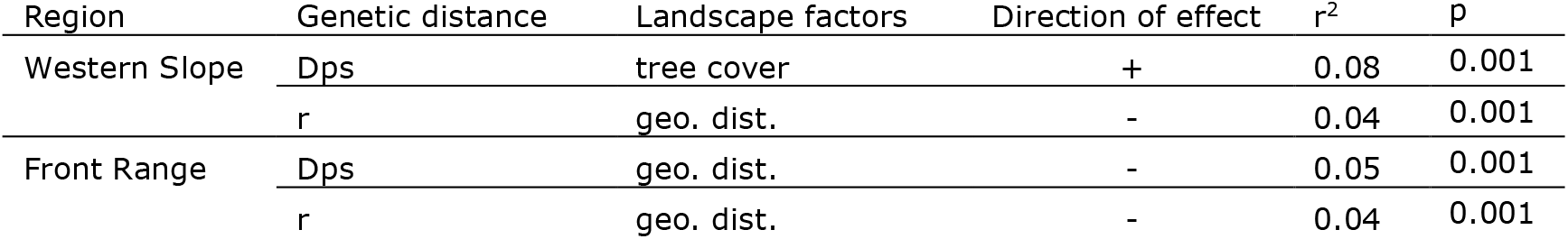
Multiple regression on distance matrices (MRDM) landscape genomic results from the Western Slope and Front Range of Colorado. Response variables were individual-based genetic distances, i.e., proportion of shared alleles (Dps) and relatedness (r). Explanatory variables, after removing correlated variables, were the geographic (Euclidean) distance model (geo. dist.), percent impervious surface cover, percent tree canopy cover, river and stream riparian corridors, roads, and minimum temperature of the coldest month. Forward selection followed by backward elimination was performed, with 1,000 random permutations of the dependent distance matrix per step, using Bonferroni-corrected p-to-enter and p-to-remove alpha values of 0.05. Standardized beta coefficients were used to assess the direction of effect of each landscape variable on gene flow. Only univariate models were supported.

**Table 5:**
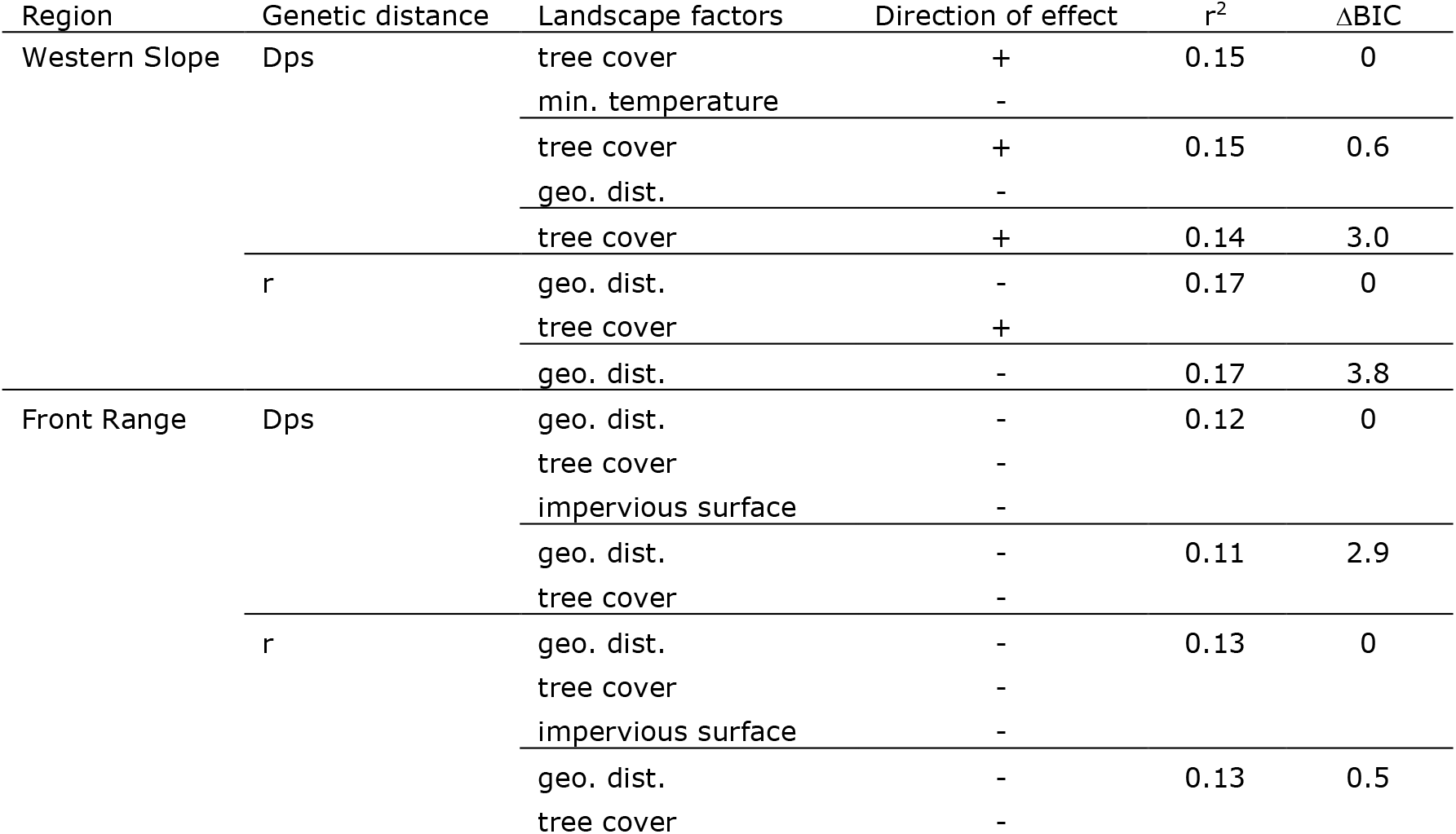
Maximum likelihood of population effects (MLPE) landscape genomic results from the Western Slope and Front Range of Colorado. Response variables were individual-based genetic distances, i.e., proportion of shared alleles (Dps) and relatedness (r). Pairwise comparisons of individuals were controlled as a random effect. Fixed effects, after removing correlated variables, were the geographic (Euclidean) distance model (geo. dist.), percent impervious surface cover, percent tree canopy cover, vegetation density, river and stream riparian corridors, roads, and minimum temperature of the coldest month. Standardized beta coefficients were used to assess the direction of effect of each landscape variable on gene flow. Models reported are within the top 5 BIC units. Landscape factors are in order of standardized beta coefficients (largest to smallest).

## Discussion

The apex predator puma (*Puma concolor*) persists in many urbanized regions throughout its range, yet the localized effects of recent urban sprawl remain unclear. Here, we compared patterns of genomic landscape connectivity and diversity of pumas across two regions that span an urban-rural divide in Colorado, USA. Landscape genomic connectivity patterns differed between regions, such that genetic distances were higher and urbanization (i.e., percent impervious surface cover) restricted gene flow in the more urbanized Front Range, whereas forest cover was most important for enhancing gene flow on the rural Western Slope. Despite finding reductions in gene flow associated with urbanization on the Front Range, population-level genetic diversity and inbreeding measures were similar to those on the rural Western Slope. This suggests that recent urban sprawl in the Colorado Front Range has not yet had a substantial impact on the genetic diversity of pumas. This is in contrast to more isolated puma populations in other highly urbanized landscapes such as southern California and Florida, which exhibit reduced genetic diversity and strong evidence of inbreeding compared to Colorado pumas (Ernest *et al.* 2003, 2014; Johnson *et al.* 2010). However, a smaller effective population size, higher among-individual genetic distances, and higher population substructure in the recently urbanized Front Range suggest habitat fragmentation has already impacted this population and could cause further reductions of genetic diversity as urbanization continues to expand in Colorado (Theobald 2005; U.S. Census Bureau 2017). If puma populations decline, this could have important cascading effects into lower trophic levels, such as overgrazing of vegetation by ungulate herbivores (Markovchik-Nicholls *et al.* 2008).

### Population genomics and structure

The Western Slope and Front Range were resolved as two genetically distinct groups (i.e., K=2; Figures 1 and 2). Minimum temperature of the coldest month was also negatively associated with gene flow in one of the top landscape genomic models on the Western Slope (Table 5), suggesting there may be restricted gene flow through high elevation, alpine tundra habitats (McMahon *et al.* 2001). However, potential immigrants and admixed individuals were identified moving in both directions (Figure 2) and overall genetic differentiation between the two populations was low (pairwise F_ST_ = 0.02; Table 2). Since our sample archive consisted of opportunistically collected samples, our analyses were restricted to populations in two distinct regions, whereas pumas occur throughout the southern Rocky Mountains in Colorado. Therefore, potential immigrants and admixed individuals are not necessarily moving between our specific Western Slope and Front Range study areas, but may originate from other unsampled populations that share genetic ancestry with our two study regions. Nevertheless, results from our study suggest pumas may be somewhat limited in dispersing across the high elevation peaks of the Continental Divide, and future studies should attempt to sample more intensively across the entire region to further investigate this trend.

We identified similar levels of genetic diversity and inbreeding between the rural Western Slope and more urbanized Front Range (Table 2), suggesting urbanization is not yet having a large impact on the genetic diversity of pumas in Colorado. One potential explanation is that urbanization in the Front Range is primarily occurring on the eastern edge of the region, possibly creating a relatively impermeable urban boundary on the eastern border, but not isolating pumas in fragments or limiting their connectivity to wildland habitat to the west (Figure 1; Lewis *et al.* 2015; Blecha *et al.* 2018). Another possibility is that many of the SNPs we sampled may not have high enough mutation rates to show a strong genomic signature of the relatively recent effects of rapid urbanization occurring in the Front Range (Haasl and Payseur 2011; Allendorf *et al.* 2013). As the human population continues to expand, future urbanization could result in more fragmented populations and reductions in genetic diversity, as has been detected in other more urbanized landscapes like southern California and Florida (Ernest *et al.* 2003, 2014; Johnson *et al.* 2010).

Despite similar geographic extents and levels of genetic diversity in the Western Slope and Front Range, mean genetic distances among individuals were higher in the urban Front Range (Table 2), suggesting that fragmentation due to urbanization may be limiting puma dispersal and gene flow. In addition, a larger effective population size (*N*_*e*_) of pumas was detected on the rural Western Slope (*N*_*e*_=69.3) compared to the urban Front Range (*N*_*e*_=40.2; Table 2), with the caveat that some assumptions of this estimator are violated in both regions (e.g., closed populations with no immigration, non-overlapping generations). The effect of non-overlapping generations on *N*_*e*_ is difficult to predict (Waples *et al.* 2016), and this assumption is expected to be violated similarly in both the Western Slope and Front Range populations. Immigration, however, is expected to downwardly bias *N*_*e*_ by creating linkage disequilibrium through a multi-locus Wahlund effect (Wahlund 1928; Waples and England 2011). Thus, it is possible that the Front Range may be showing a lower *N*_*e*_ due to having more immigrants from outside populations than the Western Slope. This is possible, and perhaps likely, given the higher overall population substructure in the Front Range (Figure 2), which could indicate more potential immigrants into this region. On the other hand, if immigration rates are similar for both regions, the relatively smaller Front Range *N*_*e*_ may be due to (1) urbanization and fragmentation impacting and limiting population size, and/or (2) species range limit theory (Abundant Center Hypothesis) predicting that smaller population sizes are likely to occur at the edge of the geographic range relative to core areas (Brown 1984; Sagarin and Gaines 2002). These potential underlying factors are not mutually exclusive and may both be acting together. However, the lack of difference in most genetic diversity measures, in addition to slightly lower allelic richness in the Front Range, which is the most sensitive metric to recent bottlenecks (Allendorf *et al.* 2013), suggests lower effective population size on the Front Range may be more consistent with recent urbanization impacts than historical range boundary effects.

### Landscape genomics

With regard to general landscape genomics methodology, we found MRDM to be a much more conservative approach that adds fewer explanatory variables to the models than MLPE (Tables 4 and 5). Conversely, MLPE results in more complex models with more explanatory variables and higher r^2^ values (genetic variation explained) than MRDM (Tables 4 and 5). The different genetic distance measures we used (D_PS_ and r) showed largely consistent relationships with landscape variables, but still provided a few different insights, particularly using MLPE (Tables 4 and 5). Overall r^2^ values were somewhat low (r^2^ = 0.04 - 0.08 for MRDM, r^2^ = 0.11 - 0.17 for MLPE), but this is expected for a large carnivore with extreme long distance dispersal abilities (e.g., Short Bull *et al*. 2011, Balkenhol *et al*. 2016). Isolation by distance was important across models for both regions (Tables 4 and 5).

On the rural Western Slope, tree canopy cover was most important for enhancing gene flow, suggesting pumas prefer to disperse through forests rather than more open shrub and grassland habitats in this landscape (Table 5). Forests provide more cover for concealment and ambush predation (Logan and Sweanor 2001; Hornocker and Negri 2009; Warren *et al.* 2016). Use of open areas may also increase susceptibility to mortality by hunters and ranchers (Newby *et al.* 2013), which are both more prevalent in the rural Western Slope than the more urbanized Front Range. In addition, non-forested areas on the Western Slope are dry, high elevation desert habitats (i.e., the Colorado Plateau ecoregion; McMahon *et al.* 2001), which may provide less prey and water resources, and thus be poorer habitats for hunting and dispersal (Sweanor *et al.* 2000; McRae *et al.* 2005; Dickson *et al.* 2013).

In the more urbanized Front Range, impervious surface cover restricted gene flow (Table 5). This suggests urbanization is limiting gene flow, despite high levels of genetic diversity (Table 2). Similarly, Lewis *et al.* (2015) found pumas were less likely to be detected in habitats with residential development, even low-density exurban developments, which are increasingly encroaching into the foothills of the Front Range region. Genetic studies on pumas from more urbanized and fragmented populations in southern California and Florida have detected strong inbreeding and isolation associated with urbanization (Ernest *et al.* 2003, 2014; Johnson *et al.* 2010; Riley *et al.* 2014). Our study detected more subtle impacts of urbanization in a less fragmented landscape, within mountainous wildland habitats adjacent to a major metropolitan center, which experiences high levels of human outdoor recreation activities such as hiking and skiing (Figure 1). In addition, in contrast with the rural Western Slope and contrary to our initial hypotheses, forest cover was negatively associated with gene flow on the Front Range (Table 5). This pattern suggests pumas are more willing to disperse through open shrub and grassland habitats in this region. The reasons for this are unclear, but pumas living in the more developed Front Range may be more acclimated to human activities and thus more willing to travel outside of forested habitats, demonstrating that pumas have a range of adaptable behaviors and will use and move through different types of habitat (Dickson *et al.* 2005; Blecha *et al.* 2018). Pumas may also be hunting more urban mesopredators, domestic, and agricultural animals in these open habitats on the more developed Front Range, which was shown in a prior study using stable isotope analysis of Front Range puma diets (Moss *et al.* 2016b). There is also less hunting of pumas in the Front Range compared to the rural Western Slope, so pumas may be less wary of open areas, although this effect would be expected to be counteracted in part by higher traffic mortality in the more urbanized region (Beier 1995; Crooks 2002).

### Conclusions

Our findings are consistent with prior comparative landscape genetic studies that have revealed varying effects of landscape factors on movement and gene flow across different portions of a species’ geographic range (e.g., Vandergast *et al.* 2007; Short Bull *et al.* 2011; Trumbo *et al.* 2013). We found that in the rural Western Slope with high hunting pressure, forests with high tree canopy cover are most important for conserving puma genetic connectivity. In contrast, in the more urbanized Front Range, non-forested habitats such as shrublands and grasslands habitats are utilized for dispersal and gene flow, effective population sizes are smaller, genetic distances among individuals are higher, and gene flow is being restricted by urbanization (Tables 2, 4, and 5). Next generation sequencing techniques can provide dense, genome-scale SNP datasets of thousands of putatively neutral markers, which gives researchers increased power to detect the often subtle effects of landscape factors, such as urbanization, on gene flow (Luikart *et al.* 2003; Lowe and Allendorf 2010; Allendorf *et al.* 2013). This is particularly important for wide-ranging species with broad geographic distributions, since landscape effects on gene flow occur at broader geographic scales and may be weaker and more difficult to detect compared to more dispersal-limited species with smaller home ranges (Holderegger *et al.* 2006; Epps *et al.* 2007; Balkenhol *et al.* 2016). Indeed prior work on pumas using 16 microsatellites found no population structure across the southern Rocky Mountains of Colorado and northern New Mexico (McRae *et al.* 2005). Our results demonstrate that large SNP datasets can allow researchers to identify impacts of urbanization on gene flow, effective population sizes, and patterns of population genetic structure of wide-ranging species, even before fragmentation is extensive enough to greatly reduce genetic diversity. Maintaining genetic connectivity in these “umbrella” species can have outsized benefits towards conserving biodiversity, since preserving broad swaths of contiguous habitats that are necessary for their persistence also benefits many other species with smaller home ranges and narrower habitat requirements (Sergio *et al.* 2006, 2008; Thorne *et al.* 2006).

## Acknowledgements

Funding was provided by the National Science Foundation, Ecology of Infectious Disease Program (NSF-EID 1413925 and 723676). Samples were collected by Colorado Parks and Wildlife. We also thank Michael Antolin, Kelly Pierce, and Jill Gerberich at Colorado State University for assistance in the lab.

## Author Contributions

D.R.T. performed laboratory work, analyzed landscape and population genomic data, and wrote the manuscript; P.S., R.B.G., C.P.K, S.K., and N.F.J. performed laboratory work and analyzed landscape and population genomic data; K.L. and M.A. directed fieldwork and collected field data; M.E.C., S.C., H.B.E., K.C., S.V., and W.C.F. conceived of study questions and directed research; and all authors contributed input to draft and final versions of the manuscript.

## Data Accessibility

ddRADseq data used in genomic analyses will be uploaded to Dryad (datadryad.org) upon acceptance of the manuscript for publication.

## Supporting Information

**Table S1**: Library replicate analysis of error rates from different Stacks parameter settings for minimum number of identical raw reads required to create a stack (−m), number of mismatches allowed between loci when processing a single individual (−m), number of mismatches allowed between loci when building the catalog (−n), and maximum number of stacks at a single de novo locus (-max_locus_stacks). Locus error rate was the number of loci present in only one of the samples of a replicate pair divided by the total number of loci, allele error rate was the number of allele mismatches between replicate pairs divided by the number of loci, and SNP error rate was the proportion of SNP mismatches between replicate pairs.

**Table S2**: Landscape resistance transformations for the Western Slope and Front Range.

**Table S3**: Correlations (Pearson’s r) between environmental raster surfaces used in landscape genomic analyses for the Western Slope and Front Range regions of Colorado. Pearson’s correlations > 0.7 are in bold.

**Table S4**: Correlations (Pearson’s r) between Circuitscape environmental resistances used in landscape genomic analyses for the Western Slope and Front Range regions of Colorado. Pearson’s correlations > 0.7 are in bold.

**Figure S1**: Principle Components Analyses (PCAs) and Admixture plots of (a) 78 Western Slope pumas and (b) 56 Front Range pumas, analyzed separately within each region.

