## Supporting Information for "Urbanization impacts apex predator gene flow but not genetic diversity across an urban-rural divide"

Table S1: Library replicate analysis of error rates from different Stacks parameter settings for minimum number of identical raw reads required to create a stack (-m), number of mismatches allowed between loci when processing a single individual (-M), number of mismatches allowed between loci when building the catalog (-n), and maximum number of stacks at a single de novo locus (-max_locus_stacks). Locus error rate was the number of loci present in only one of the samples of a replicate pair divided by the total number of loci, allele error rate was the number of allele mismatches between replicate pairs divided by the number of loci, and SNP error rate was the proportion of SNP mismatches between replicate pairs. Final parameter settings are highlighted in bold.

| Stacks parameter settings | | | | Error rates | | | Total number of SNPs |
|---|---|---|---|---|---|---|---|
| -m | -M | -n | -max_locus_stacks | Loci error | Allele error | SNP error | |
| 3 | 2 | 2 | **3** | 0.217 | 0.129 | 0.0758 | 29,448 |
| 3 | 2 | 2 | 4 | 0.217 | 0.129 | 0.0761 | 29.434 |
| 3 | 2 | 2 | 5 | 0.212 | 0.129 | 0.0761 | 29,412 |
| 3 | 2 | **4** | 3 | 0.217 | 0.128 | 0.0740 | 29,814 |
| 3 | 2 | 6 | 3 | 0.216 | 0.128 | 0.0732 | 28,984 |
| 3 | 2 | 8 | 3 | 0.216 | 0.128 | 0.0730 | 28,507 |
| 3 | **4** | 2 | 3 | 0.217 | 0.128 | 0.0772 | 30,658 |
| 3 | 6 | 2 | 3 | 0.216 | 0.127 | 0.0763 | 29,636 |
| **3** | 8 | 2 | 3 | 0.217 | 0.127 | 0.0756 | 28,796 |
| 5 | 2 | 2 | 3 | 0.217 | 0.129 | 0.0758 | 23,062 |
| 7 | 2 | 2 | 3 | 0.219 | 0.065 | 0.0582 | 18,541 |

Table S2: Landscape resistance transformations for the Western Slope and Front Range.

| Landscape Resistances | Western Slope | | | | | Front Range | | | | | Transformations |
|---|---|---|---|---|---|---|---|---|---|---|---|
| | Min | Max | Median | Mean | Std Dev | Min | Max | Median | Mean | Std Dev | |
| Elevation (m) | 1 | 2910 | 2009 | 1974.4 | 529.3 | 1 | 2874 | 1984 | 1973.6 | 629.3 | Convert to integer, subtract minimum value, add 1. |
| Tree canopy cover (%) | 1 | 101 | 81 | 71.1 | 31.4 | 1 | 101 | 69 | 66.0 | 33.6 | Add 1, invert raster. |
| Impervious surface (%) | 1 | 101 | 1 | 1.5 | 4.1 | 1 | 101 | 1 | 5.0 | 13.5 | Add 1. |
| Minimum temp. coldest month (°C) | 1 | 108 | 39 | 44.5 | 28.1 | 1 | 117 | 45 | 44.0 | 28.0 | Multiply by 10, add minimum value, add 1, invert raster. |
| Annual precipitation (mm) | 1 | 930 | 680 | 654.7 | 171.5 | 1 | 648 | 555 | 510.6 | 121.9 | Subtract minimum value, add 1, invert raster. |
| Enhanced vegetation index | 1 | 10814 | 4374 | 4573.1 | 1858.3 | 1 | 11102 | 3717 | 4175.2 | 1800.9 | Add minimum value, add 1, invert raster. |
| Topographic roughness | 1 | 27925 | 11 | 50.5 | 129.3 | 1 | 20068 | 20043 | 20010.7 | 101.8 | Convert to integer, add 1, invert raster. |
| Landcover | 1 | 10 | 1 | 3.2 | 3.0 | 1 | 10 | 1 | 4.2 | 3.7 | Forested habitats = 1, natural, non-forested habitats (e.g., grasslands, shrublands) = 5, human developed, barren, and agricultural areas = 10. |
| Roads | 1 | 10 | 1 | 1.7 | 2.5 | 1 | 10 | 1 | 2.7 | 3.6 | Roads = 10, non-roads = 1. |
| Riparian | 1 | 10 | 10 | 9.4 | 2.3 | 1 | 10 | 10 | 9.4 | 2.3 | Riparian = 1, non-riparian = 10. |
| Raw Landscape Data | Western Slope | | | | | Front Range | | | | | Hypotheses** |
| | Min | Max | Median | Mean | Std Dev | Min | Max | Median | Mean | Std Dev | |
| Elevation (m) | 1453.5 | 4362.9 | 2354 | 2418.0 | 552.5 | 1474.5 | 4347.1 | 2365 | 2374.9 | 629.3 | Higher elevation has higher resistance to gene flow. |
| Tree canopy cover (%) | 0 | 100 | 20 | 29.9 | 31.4 | 0 | 100 | 32 | 35.0 | 33.6 | Higher tree cover has lower resistance to gene flow. |
| Impervious surface (%) | 0 | 100 | 0 | 0.5 | 4.1 | 0 | 100 | 0 | 4.0 | 13.5 | Higher impervious surface cover has higher resistance to gene flow. |
| Minimum temp. coldest month (°C) | -20.2 | -9.5 | -13.3 | -13.9 | 2.8 | -19.9 | -8.3 | -12.7 | -12.6 | 2.8 | Higher minimum temperature has lower resistance to gene flow. |
| Annual precipitation (mm) | 208 | 1137 | 458 | 483.3 | 171.5 | 359 | 1006 | 452 | 496.4 | 121.9 | Higher precipitation has lower resistance to gene flow. |
| Enhanced vegetation index | -1806 | 8955 | 4634 | 4434.9 | 1858.3 | -1969 | 9132 | 5416 | 4957.8 | 1800.9 | Higher vegetation density has lower resistance to gene flow. |

| | | | | | | | | | | | |
|---|---|---|---|---|---|---|---|---|---|---|---|
| Topographic roughness | 0 | 27924.6 | 11 | 53.1 | 129.6 | 0 | 20067.0 | 25 | 56.2 | 100.8 | Higher topographic roughness has lower resistance to gene flow. |
| Landcover* | N/A | N/A | N/A | N/A | N/A | N/A | N/A | N/A | N/A | N/A | Forested habitats have lowest resistance; natural, non-forested habitats (e.g., grasslands, shrublands) have intermediate resistance; and human developed, barren, and agricultural areas have highest resistance to gene flow. |
| Roads* | N/A | N/A | N/A | N/A | N/A | N/A | N/A | N/A | N/A | N/A | Roads have high resistance to gene flow. |
| Riparian* | N/A | N/A | N/A | N/A | N/A | N/A | N/A | N/A | N/A | N/A | Riparian corridors have low resistance to gene flow. |

*Categorical variables; no min, max, median, mean, or std dev
** Ecological justifications for each hypothesis are described in Table 1.

Table S3: Correlations (Pearson's r) between environmental raster surfaces used in landscape genomic analyses for the Western Slope and Front Range regions of Colorado. Pearson's correlations > 0.7 are in bold.

| Western Slope | Land cover | Impervious | Roads | Riparian | Tree cover | Veg. density | Min. temp. | Ann. precip. | Topo. rough. | Elevation |
|---|---|---|---|---|---|---|---|---|---|---|
| Land cover | 1 | -0.18 | -0.12 | 0.02 | 0.64 | 0.30 | -0.15 | 0.11 | -0.04 | 0.14 |
| Impervious | | 1 | 0.31 | 0.01 | -0.08 | -0.01 | 0.09 | -0.10 | -0.04 | -0.10 |
| Roads | | | 1 | 0.02 | -0.08 | 0.04 | 0.06 | -0.09 | -0.09 | -0.09 |
| Riparian | | | | 1 | 0.01 | 0.04 | 0.05 | -0.06 | -0.02 | -0.08 |
| Tree cover | | | | | 1 | 0.32 | -0.29 | 0.25 | 0.08 | 0.28 |
| Veg. density | | | | | | 1 | -0.01 | -0.16 | -0.19 | -0.10 |
| Min. temp. | | | | | | | 1 | **-0.81** | -0.27 | **-0.88** |
| Ann. precip. | | | | | | | | 1 | 0.27 | **0.98** |
| Topo. rough. | | | | | | | | | 1 | 0.28 |
| Elevation | | | | | | | | | | 1 |

| Front Range | Land cover | Impervious | Roads | Riparian | Tree cover | Veg. density | Min. temp. | Ann. precip. | Topo. rough. | Elevation |
|---|---|---|---|---|---|---|---|---|---|---|
| Land cover | 1 | -0.45 | -0.32 | -0.03 | **0.79** | 0.15 | -0.37 | 0.15 | 0.05 | 0.40 |
| Impervious | | 1 | 0.47 | 0.01 | -0.27 | -0.01 | 0.33 | -0.21 | -0.12 | -0.32 |
| Roads | | | 1 | 0.07 | -0.23 | 0.09 | 0.30 | -0.24 | -0.15 | -0.31 |
| Riparian | | | | 1 | -0.04 | 0.05 | 0.09 | -0.08 | -0.04 | -0.11 |
| Tree cover | | | | | 1 | 0.13 | -0.39 | 0.19 | 0.05 | 0.41 |
| Veg. density | | | | | | 1 | 0.55 | **-0.71** | -0.21 | -0.60 |
| Min. temp. | | | | | | | 1 | **-0.86** | -0.23 | **-0.95** |
| Ann. precip. | | | | | | | | 1 | 0.25 | **0.91** |
| Topo. rough. | | | | | | | | | 1 | 0.26 |
| Elevation | | | | | | | | | | 1 |

Table S4: Correlations (Pearson's r) between Circuitscape environmental resistances used in landscape genomic analyses for the Western Slope and Front Range regions of Colorado. Pearson's correlations > 0.7 are in bold.

Western Slope

| | Null | Land cover | Impervious | Roads | Riparian | Tree cover | Veg. density | Min. temp. | Ann. precip. | Topo. rough. | Elevation |
|---|---|---|---|---|---|---|---|---|---|---|---|
| Null | 1 | 0.60 | **0.83** | 0.44 | 0.67 | 0.68 | **0.76** | 0.61 | **0.95** | **0.99** | **0.95** |
| Land cover | | 1 | 0.63 | 0.31 | 0.45 | **0.80** | 0.68 | 0.35 | 0.62 | 0.60 | 0.61 |
| Impervious | | | 1 | 0.62 | 0.50 | 0.60 | 0.63 | 0.49 | **0.81** | **0.83** | **0.81** |
| Roads | | | | 1 | 0.28 | 0.27 | 0.38 | 0.28 | 0.43 | 0.44 | 0.43 |
| Riparian | | | | | 1 | 0.45 | 0.53 | 0.54 | 0.58 | 0.67 | 0.57 |
| Tree cover | | | | | | 1 | **0.72** | 0.19 | **0.73** | 0.68 | **0.72** |
| Veg. density | | | | | | | 1 | 0.37 | **0.80** | **0.76** | **0.78** |
| Min. temp. | | | | | | | | 1 | 0.38 | 0.61 | 0.38 |
| Ann precip. | | | | | | | | | 1 | **0.95** | **0.99** |
| Topo. rough. | | | | | | | | | | 1 | **0.95** |
| Elevation | | | | | | | | | | | 1 |

Front Range

| | Null | Land cover | Impervious | Roads | Riparian | Tree cover | Veg. density | Min. temp. | Ann. precip. | Topo. rough. | Elevation |
|---|---|---|---|---|---|---|---|---|---|---|---|
| Null | 1 | 0.30 | 0.10 | 0.43 | 0.65 | 0.62 | **0.91** | 0.50 | **0.98** | **0.99** | **0.89** |
| Land cover | | 1 | **0.87** | **0.71** | -0.04 | **0.75** | 0.32 | -0.19 | 0.33 | 0.30 | 0.47 |
| Impervious | | | 1 | 0.66 | -0.16 | 0.47 | 0.14 | -0.22 | 0.13 | 0.10 | 0.26 |
| Roads | | | | 1 | 0.17 | 0.60 | 0.43 | 0.15 | 0.42 | 0.43 | 0.44 |
| Riparian | | | | | 1 | 0.19 | 0.50 | 0.61 | 0.58 | 0.65 | 0.44 |
| Tree cover | | | | | | 1 | 0.63 | 0.04 | 0.65 | 0.63 | **0.73** |
| Veg. density | | | | | | | 1 | 0.47 | **0.89** | **0.90** | **0.76** |
| Min. temp. | | | | | | | | 1 | 0.35 | 0.50 | 0.08 |
| Ann. precip. | | | | | | | | | 1 | **0.98** | **0.94** |
| Topo. rough. | | | | | | | | | | 1 | **0.89** |
| Elevation | | | | | | | | | | | 1 |

Table S5: Effective population sizes ($N_e$) based on subsetting individuals into different time periods.

| Western Slope | 2005-Dec.2010 | Jan.2011-2014 |
|---|---|---|
| $N_{gen}$ | 35 indiv. | 43 indiv. |
| $N_e$ (95% C.I.) | 38.5 (37.0-40.4) | 82.2 (79.0-85.5) |
| Front Range | 2007-May2010 | June2010-2013 |
| $N_{gen}$ | 25 indiv. | 21 indiv. |
| $N_e$ (95% C.I.) | 34.4 (33.0-35.8) | 35.2 (33.8-36.6) |

Figure S1: Principle Components Analyses (PCAs) and Admixture plots of (a) 78 Western Slope pumas and (b) 56 Front Range pumas, analyzed separately within each region.

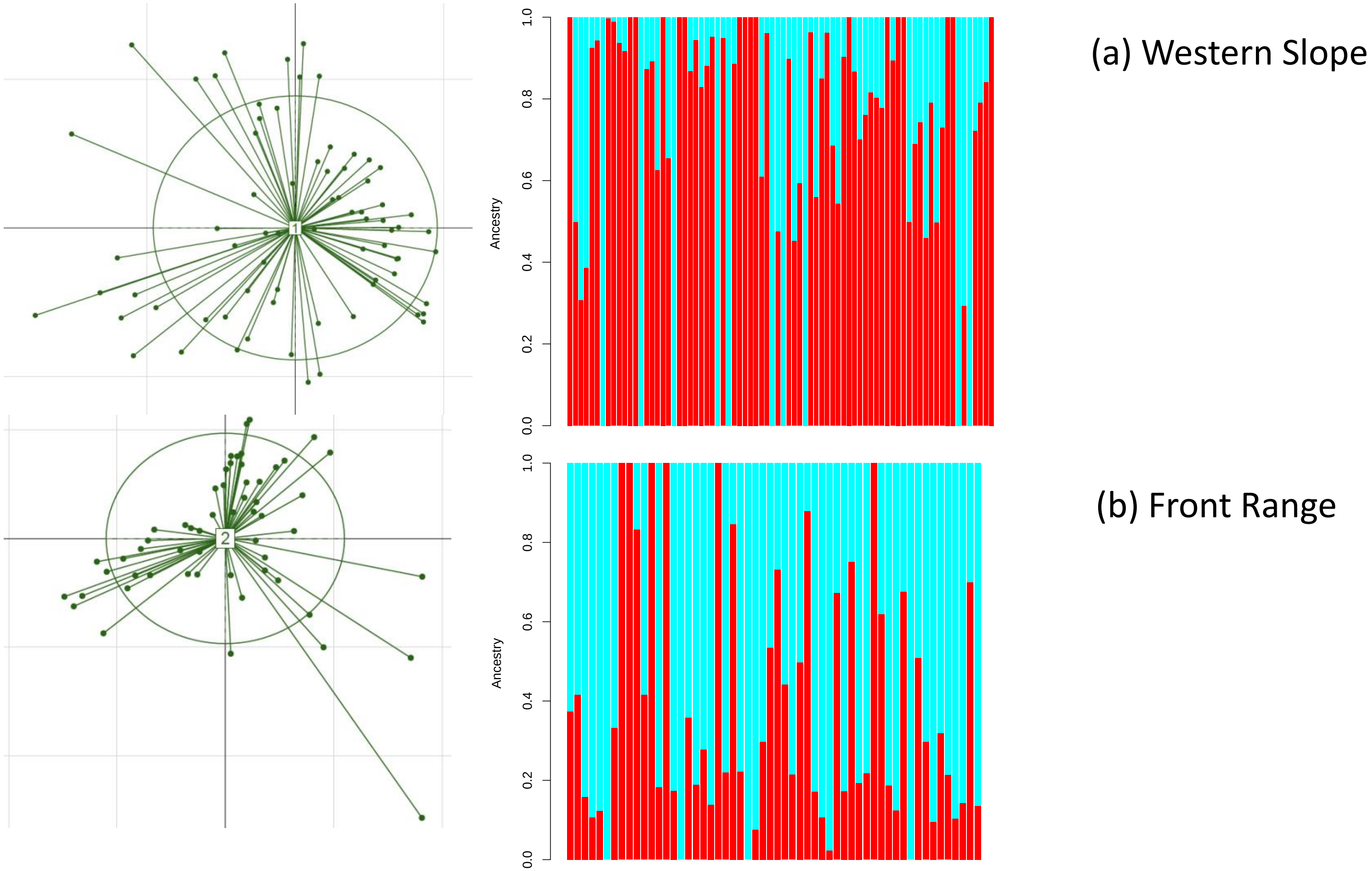
